# Long-term Nrf2-driven microglial repopulation mitigates microgliosis, neuronal loss and cognitive deficits in tauopathy

**DOI:** 10.1101/2025.06.11.659019

**Authors:** Lucía Viqueira, Elisa Navarro, Pilar Negredo, Juan Antonio Bernal, María Isabel Rodríguez-Franco, Elena Tortosa, Manuela G. López

## Abstract

A prominent pathological feature of tauopathies, including Alzheimer’s disease, is a chronic microglial reactivity, which contributes to neuroinflammation and disease progression. Microglia, the innate immune cells of the brain, can be pharmacologically eliminated by inhibiting the colony stimulating factor 1 receptor (e.g. with PLX5622), which is essential for their survival and proliferation. Upon inhibitor withdrawal, microglia rapidly repopulate, replenishing the central nervous system niche with naïve cells. In recent years, microglia repopulation strategies have gained great interest as a means to reprogram dysfunctional microglia, while avoiding the detrimental effects of prolonged immune depletion. Despite promising short-term results, the long-term efficacy and pharmacological modulation of repopulated microglial remain poorly understood. Here, we investigate whether repopulated microglia after PLX5622 treatment sustains their beneficial effects over time in an AAV-hTauP301L induced model. Additionally, we assessed whether activating the cytoprotective nuclear factor erythroid 2 p45–related factor 2 (Nrf2) during microglial repopulation enhanced and prolonged the therapeutic outcomes. While microglial repopulation alone failed to maintain its neuroprotection in the long-term, when combined with an Nrf2 inducer, it improved cognitive deficits, reverted hippocampal neuronal loss and restored microglial phenotypes and mitochondrial energetics homeostasis, in our tauopathy-induced model. These results highlight the importance of shaping the fate of self-renewed microglia and propose Nrf2-mediated microglia repopulation as a potential pharmacological strategy for the treatment of tauopathies.

**HIGHLIGHTS:**

- Nrf2 inducer ND523 boosts long-term repopulated microglia function in tauopathy.
- ND523-mediated microglial repopulation mitigates tau-induced cognitive decline.
- ND523-driven microglial repopulation reduces neuronal loss without altering p-tau.
- ND523-microglial repopulation rescues microglial morphology and reduces reactivity.
- ND523-mediated repopulation prevents mitochondrial energetics dysfunctions.

## 1. INTRODUCTION

Alzheimer’s disease (AD) is the leading cause of dementia worldwide, representing a major health care challenge in a world with demographic ageing. AD belongs to a heterogenous group of neurodegenerative disorders known as tauopathies, characterized by abnormal accumulation of tau protein aggregates within neurons ^1^. Tau is a microtubule-associated protein which plays a critical role in the assembly and stabilization of microtubules under normal conditions. However, in tauopathies, it undergoes a series of post-translational modifications, that lead to misfolding, aggregation and intracellular accumulation as neurofibrillary tangles (NFTs), a hallmark of these diseases ^2^. Notably, AD is considered a secondary tauopathy due to the concomitant presence of amyloid-beta (Aβ) plaques, which precede tau pathology ^2,3^. Beyond these two main histopathological hallmarks, increasing evidence highlights the significant role of the inflammatory component ^4–6^, redefining AD not simply as a neuron-focused disease but as one profoundly shaped by the brain’s immune landscape ^7^.

Microglia, the primary immune cells of the central nervous system (CNS), account for approximately 10 % of the cell population in the brain. They are highly dynamic ramified cells, continuously surveying the local brain environment via their processes and phagocyting pathogens and debris ^8^. They also participate in a broader array of CNS functions; from vasculo-, glio-, and neurogenesis, to myelination or synapse remodeling ^9^. In response to brain damage or disease, microglia undergo morphological, transcriptional, and functional changes as an initial beneficial response ^10^. If unresolved, this chronic reactive state of the brain’s immune system drives neuronal damage and exacerbates disease progression ^11^.

Therapeutically, targeting microglia to modulate their number and function has gained significant interest. One promising approach involves the pharmacological inhibition of the colony-stimulating factor 1 receptor (CSF1R), a crucial signaling pathway for microglial survival and proliferation. Among available inhibitors, PLX5622 stands out for its efficacy, specificity and brain penetrance ^12^, reaching a near-complete microglial depletion in murine models ^13–16^. Upon inhibitor removal, residual microglia rapidly replenish the emptied CNS niche within a few days ^17,18^. Depletion strategies have been used to study the effects of microglia absence in AD hallmark disease features. In tauopathy mouse models, microglial ablation halts pathology progression ^19^, suppresses tau propagation ^20^ and attenuates tau-induced neurodegeneration ^21^. Strikingly, in the Aβ context, early preventive treatments with CSF1R inhibitors reduce plaque formation ^14,22^, whereas in mice with established brain Aβ plaques, microglial depletion strategies do not modify Aβ levels or plaque load while they prevent cognitive deficit ^15,23^, dendritic spine and neuronal loss ^16^.

Prolonged microglial elimination is unlikely to be clinically feasible due to potential risks associated with chronic depletion of the braińs immune system. As a result, in recent years, microglial repopulation strategies have emerged, aiming to replace dysfunctional microglia with newly generated functional cells. Studies have shown that this approach not only improves cognition but also restores microglial phenotypes in aged mice ^24^ and in amyloid-β–driven models, such as the 5xFAD mouse ^25^. However, most of these investigations have been confined to the Aβ context, leaving a significant gap in our understanding of repopulated microglia in tau-driven models. In addition, they have been limited to short-term observations, leaving unclear whether these newly introduced cells can maintain their homeostatic, non-reactive phenotype or whether they may gradually revert to the reactive state they displayed prior to depletion. A promising target to support long-term microglial function is the nuclear factor Nrf2, a master regulator of cellular stress response. Nrf2 modulates the expression of numerous cytoprotective genes involved in antioxidant and anti-inflammatory responses ^26^, which could have a positive impact in preserving microglial function over time.

Herein, we have evaluated the potential benefits of microglial repopulation over a prolonged period in a tau-driven model. We also evaluated the ability of compound ND523, a novel Nrf2 activator developed by our group ^27^, to enhance the efficacy of microglia repopulation. Our work provides critical insights into the limitations of self-renewed microglia in maintaining their functions over time and highlights the potential benefits of the combined therapy using a CSF1R inhibitor followed by an Nrf2 inducer to reshape the microglial landscape over the long term. In doing so, we contribute to a growing body of evidence that positions microglia and its modulation as central therapeutic targets in neurodegenerative diseases, and more specifically to tauopathies.

## 2. MATERIALS AND METHODS

### 2.1. Animal use and care

Male C57BL/6J wild-type (WT) mice 4-5 months old were used in all experiments. Only males were selected to ensure effective depletion, as observed in previous studies ^19,28^. Mice were originally obtained from Charles River Laboratories and bred in-house. Animals were housed in a conventional animal facility at the Autonomous University of Madrid (Madrid, Spain) under controlled humidity and temperature conditions, with a 12-hour light/dark cycle. Rodent chow and water were provided *ad libitum*, and cages with bedding were changed weekly. All experiments were conducted in accordance with the European Union Directive 2010/63/EU on the protection of animals used for experimental and other scientific purposes and the Spanish Royal Decree for Animal Protection RD53/2013. The study protocol was approved by the Institutional Ethics Committee of the Autonomous University of Madrid and the Community of Madrid, Spain (PROEX: 218.5/20). Efforts were made to minimize animal use and reduce suffering at all stages of the study.

### 2.2. Adeno-associated viral vectors AAV-Syn1-GFP/AAV-Syn1-hTau^P301L^ generation

Recombinant adeno-associated viral (AAV) vectors with the serotype 6 were produced and purified as previously described ^29^ with slight modifications. Plasmids were generated and propagated in *Escherichia coli* strain Stbl3 (Life Technologies). Human embryonic kidney (HEK293T) cells were transiently double-transfected using polyethylenimine (Polysciences) with the plasmids: the genome plasmid containing AAV-2 ITRs and the human promoter for synapsin-1 (SYN-1) gene limiting the expression of either green fluorescent protein (GFP) or human Tau P301L mutation and the helper plasmid pDP6 (Plasmid Factory) containing the helper sequences plus the Rep gene from AAV2 and the Cap gene from AAV6. Seventy-two hours post-transfection, HEK293T cells were centrifuged, and the resulting pellet was resuspended in lysis buffer (150 mM NaCl, 50 mM Tris-HCl pH= 8) and subjected to three freeze/thaw cycles. Then, the suspension was treated with 150 U/mL of Benzonase (Millipore) at 37°C for 30 minutes. Purification of the AAV vectors was performed using iodixanol step gradient ultracentrifugation, followed by concentration with Amicon UltraCel columns (Millipore). Viral titters were quantified via quantitative PCR (qPCR), and purified AAVs were stored at -80°C until further use.

### 2.3. Tauopathy mouse model induced by stereotaxic injection of AAV-hTau^P301L^

Tauopathy was induced by bilateral intrahippocampal stereotaxic injection of recombinant AAV vectors expressing pathological human tau mutation P301L (AAV-hTau^P301L^), under the control of the neuron-specific promoter SYN-1, limiting its expression to neurons. Control animals received injections of similar AAV particles but expressing GFP (AAV-GFP). Surgeries were conducted under Isoflutek (isoflurane 1,000 mg/mL) anesthesia in oxygen, with continuous monitoring and regulation of the animal’s temperature using a rectal probe and heating pad (Cibertec). Stereotaxic injections were performed on adult mice using a stereotaxic frame (Stoelting, Wood Dale, IL, USA). Animals’ heads were shaved, and the skin was disinfected with povidone-iodine (Betadine) prior to incision. Using a micro-drill, two cranial perforations were made bilaterally at coordinates at 1.94 mm anterior-posterior and ± 1.4 medial-lateral to bregma. A 10 µl Hamilton syringe (1701 RN Neuros Syringe) coupled to an injector (KDS Legacy) was used to bilaterally deliver a volume of 1.01 µL of AAVs (4.39·10^12^ VP/mL-1) with a rate of 0.15 µL/minute, into each hippocampus at -1.8 mm dorsal-ventral. Following the injection, the skin was stitched back, and the animal was housed for postoperative monitoring. Animals were randomly assigned into different experimental groups.

### 2.4. Animal treatments

Plexxikon Inc. supplied PLX5622 compound and Research Diets Inc formulated it in AIN-76A standard chow at 1200 ppm. To ensure microglial depletion, PLX5622 was administered for 14 days, from 7 days-post injection (dpi) of AAV until 21 dpi. Afterwards, PLX5622 chow was withdrawn to stimulate microglial repopulation.

ND523, [(*R*,*S*)-5-methoxy-3-(5-methoxyindolin-2-yl)-1*H*-indole], administration was formulated in gelatin. For this purpose, a dose of 50 mg/kg was embedded into a gelatine mixture composed of 10 % porcine skin gelatine (Sigma-Aldrich) and 10 % sucralose (DulciLight) in hot water. The mixture was gently stirred at room temperature (RT) until it cooled, and the compound was added. Once thoroughly mixed, 1 mL of the formulation was dispensed into each individual cavity of a mould. Once solidified, the jellies were unmolded and fed mice. Mice in the ND523-treated groups received a daily ND523-jelly dose from 21 dpi until the study endpoint, while the other groups received control jellies (without the compound). To ensure jelly acceptance, all mice (regardless of group) were trained with control jellies prior to beginning of the actual treatment.

### 2.5. Locomotor activity assessment

The Open Field test was used to evaluate the locomotor activity of mice, as reported in ^30^. Briefly, mice were individually placed within a 40-cm-side square box and recorded while moving freely and exploring the environment. Mice movement was tracked for a period of 5 minutes using AnyMaze software version 4.99 (Stoelting, Wood Dale, IL, USA). The 40-cm-side square box was digitally subdivided into a 4 x 4 grid (generating 16 identical squares), and different locomotor parameters were calculated, such as total travelled distance, average speed, and time/distance in the inner/outer area of the box.

### 2.6. Behavioral tests

<u>Novel Object Recognition (NOR)</u> was used to evaluate mice recognition memory as outlined in the protocol ^31^. It consists of three consecutive days (T_0_, T_1_ and T_2_), each performed 24 hours apart. During T_0_, habituation day, mice were individually located into a 40-cm-side square box and left to explore freely for 8 minutes. On the training day (T_1_), mice were placed back into the same box with two identical objects and allowed to explore for another 8 minutes. Finally, on the testing day (T_2_), one of the objects was substituted by a distinctly different object in terms of color, morphology and texture, allowing exploration of these two contrasting objects for 8 minutes. All three trials were recorded and T_2_ was used to calculate the discrimination index (DI) as an indicator of the cognitive state of the mice, being the DI equal to time spent exploring the new object minus the time spent in the old object divided by the total exploring time. Mice with no cognitive impairment will spend most of their time exploring the novel object over the familiar one, driven by their innate preference for novelty. Mice with cognitive impairment will fail to remember the object and will explore both objects equally, giving a DI close to 0.

<u>Object Location Test (OLT)</u> was used to assess spatial memory with adaptations to the method outlined in ^32^. This test spans along three consecutive days (T_0_, T_1_ and T_2_), with each trial conducted 24 hours apart. Different coloured and shaped tracks were placed on the walls of the box for orientation purposes throughout the whole test. The habituation day (T_0_) and the training day (T_1_) were conducted identically to the NOR. On the final test day (T_2_), one of the identical objects was relocated to the opposite side of the box, thus changing its location. All three trials were recorded and T_2_ was used to calculate the DI as previously mentioned, now evaluating novel *vs*. familiar location. Cognitively unimpaired mice will explore the novel location more than the familiar one. However, cognitively impaired mice fail to remember the previous object location and explore both objects equally, resulting in a DI close to 0.

Both OLT and NOR tests were performed and analysed blinded using stopwatches. The same mouse was never subjected to perform both tests.

### 2.7. Immunofluorescence analysis

42- or 84-dpi, mice were anaesthetised with Isoflutek and transcardiac perfusion was performed with ice-cold phosphate-buffered saline (PBS), followed by 4 % paraformaldehyde (PFA). Brains were extracted and post-fixed by immersion in the same fixative at 4°C overnight in mild agitation. Then, they were embedded in 30 % sucrose in 0.1 M phosphate buffer (PB) for two days before being cut into coronal sections of 40 μm on a sliding microtome. Immunostaining assays were performed on free-floating brain sections in continuous mild agitation. Initially, sections were washed extensively with PBS and permeabilized with 2 % Triton X-100 in PBS for 1 hour at RT. Then, the slices were incubated with blocking solution (10 % goat-serum in PBS 1X - 1% Triton X 100) for 2 hours at RT, followed by the primary antibodies (Table 1) incubation in blocking solution at 4 °C overnight. Next, three washing steps with PBS 1X - 0.1% Triton were performed, and the slices were incubated with the corresponding secondary antibodies (Table 2) diluted in blocking solution for 2 hours at RT. Then, three washes with PBS 1X were performed including DAPI (Thermo Fisher) (1:1000) in the second wash. Finally, slices were mounted on slides and coverslipped using Mowiol (Sigma-Aldrich) as mounting media. Images were acquired with a confocal microscope Leica TCS-SP5 (Leica Microsystems) and analysed using Fiji Software (NIH, 2.14.0/1.54h)^33^.

**Table 1.** List of primary antibodies used for immunofluorescence analysis.

| Target | Species | Dilution | Reference | Provider |
| --- | --- | --- | --- | --- |
| **AT8** | Mouse | 1:500 | MN1020 | Thermo Fisher |
| **Cd68** | Rat | 1:500 | MCA1957GA | Bio-rad |
| **C3** | Rat | 1:50 | HM1045 | Deltaclon |
| **GFAP** | Mouse | 1:1000 | MAB3402 | Sigma-Aldrich |
| **Iba 1** | Rabbit | 1:800 | 019-19741 | Wako |
| **PU.1** | Rabbit | 1:100 | 2266S | Cell Signalling |
| **PHF1** | Mouse | 1:1500 | <sup>34</sup> | Dr. Peter Davies (USA) |

**Table 2.** List of secondary antibodies used for immunofluorescence analysis.

| Excitation<br>Laser Line<br>(nm) | Species | Dilution | Reference | Provider |
| --- | --- | --- | --- | --- |
| 546 | Rat | 1:1000 | A-11081 | Thermo Fisher |
| 550 (Cyanine<br>Cy™3 ) | Mouse | 1:500 | AB_2338685 | Jackson<br>Immunoresearch |
| 550 (Cyanine<br>Cy™3 ) | Rabbit | 1:500 | AB_2338003 | Jackson<br>Immunoresearch |
| 647 | Mouse | 1:1000 | A-21237 | Thermo Fisher |
| 647 | Rat | 1:500 | A21247 | Thermo Fisher |

### 2.8. Image analysis

All image analyses were performed using Fiji software (NIH, 2.14.0/1.54h) ^33^. All experiments were carried out in hippocampal sections from similar brain coordinates. <u>Dentate Gyrus (DG) granule layer thickness</u> measurements were performed drawing a perpendicular line to the cell layer to measure the thickness. Ten different measurements were acquired per mouse and represented as mean thickness. For the analysis of <u>cluster</u> <u>of differentiation 68 (Cd68) within ionized calcium-binding adaptor molecule 1 (Iba1) and</u> <u>complement 3 (C3) within glial fibrillary acidic protein (GFAP) positive cells</u>, a mask of the area occupied by Iba1 or GFAP staining was created, and the mean intensity of Cd68 or C3 within that region was measured. In this manner, measurements were restricted to microglia or astrocytes, respectively. For <u>AT8 and PHF1</u> analysis different ROIs/zones were selected, and their mean intensity measured. Iba1^+^ cell numbers were manually counted on maximum intensity projections on the region of interest. For <u>PU.1+ cell count</u>, images were thresholded, and the number of positive nuclei was quantified using the “Analyze particles” function in a standardized macro, allowing for automated and unbiased cell counting across defined regions of interest. <u>Microglial segmentation analysis</u> was performed on Iba1 images as described previously ^35^. Briefly, images were converted to 8-bit grayscale, background-subtracted and Gaussian-blurred. An ImageJ macro from the original paper was custom and used to segment cell bodies via the “Area Maxima local maximum detection” function. Thresholds were defined as Th1 = Imax – f1 * ISD (f1 empirically set), with an area threshold of 250 px. Seed points were used for watershed analysis with Th2 = Imean + f2 * ISD (f2 set empirically). Segmented cells >1000 px were analyzed using MorphoLibJ, and different morphology readouts (area, perimeter, ramification index, cell size, Feret diameter and geodesic diameter) were obtained and analyzed.

### 2.9. RNA-sequencing sample isolation

Total RNA was extracted from hippocampal tissue using the RNeasy Mini Kit (Qiagen) following the manufacturer’s protocol, with minor adjustments. Tissue samples were lysed in 700 μL of QIAzol Lysis Reagent (Qiagen) and homogenized using a TissueLyser (Qiagen) with a 3 mm steel-ball at 50 Hz for 5 minutes (2 cycles). Subsequently, 140 μL of chloroform were added, and the samples were vortexed and centrifuged at 12,000 × g for 15 minutes at 4 °C. The upper aqueous phase was carefully transferred to a new tube. An equal volume of 70 % ethanol (600 μL) was added, and the mixture was then loaded onto RNeasy Mini spin columns using the QIAcube automated nucleic acid extraction system (Qiagen). RNA purification proceeded according to the manufacturer’s instructions. RNA concentration and purity (A260/A280, A260/A230 ratios) were assessed using a NanoDrop 8000 spectrophotometer (Thermo Fisher), and RNA integrity was evaluated with an Agilent 2100 Bioanalyzer (Agilent Technologies).

### 2.10. RNA-sequencing data generation and processing

RNA sequencing (RNA-seq) libraries were prepared from total RNA using poly-T oligo-attached magnetic beads for mRNA enrichment. Following fragmentation, cDNA synthesis was performed using random hexamer primers. Library construction included end repair, A-tailing, adapter ligation, size selection, PCR amplification, and purification. Library quality and concentration were assessed using Qubit, RT-PCR, and bioanalyzer. Libraries were pooled based on effective concentration and desired sequencing depth and sequencing was carried out on an Illumina platform using the Sequencing by Synthesis (SBS) method. Raw sequencing reads were processed using fastp software to remove adapter sequences, poly-N stretches, and low-quality reads, yielding high-quality clean reads. Q20, Q30, and GC content quality metrics were also calculated during this step. Clean reads were aligned to the GRCm38 genome using Hisat2 (v2.0.5). Genome and gene annotation files were obtained from public databases. HISAT2 was chosen for its ability to build splice-aware indices using gene annotations, improving alignment accuracy. Gene-level quantification was performed using featureCounts (v1.5.0-p3). All RNA-seq procedures described above were performed by Novogene GmbH (Munich, Germany).

### 2.11. Differential expression and pathway enrichment analysis

Expression levels were calculated as counts per million (CPM) using the cpm function from the edgeR package in R ^36^. Genes with low expression—defined as having fewer than one CPM—were filtered out, resulting in a final set of 14,013 genes. Principal Component Analysis (PCA) was performed to identify major sources of variation and detect potential outliers. To minimize technical noise arising from sequencing batch and CHIP effects, these variables were regressed out. The final dataset included 14 samples: 5 Ctrl, 4 hTau, and 5 ND-Repop. Differential expression analysis was conducted using the limma package in R ^37^. Normalization of raw count data was carried out with the edgeR package using the trimmed mean of M-values (TMM), followed by voom transformation. P-values were adjusted for multiple testing using the Benjamini-Hochberg false discovery rate (FDR) method. Gene Set Enrichment Analysis (GSEA) was performed, focusing on Hallmark gene sets ^38^, using the fGSEA package in R ^39^. Ranked differentially expressed genes (DEGs), based on t-values, were used as input. Only pathways with an adjusted P-value < 0.05 were considered significant.

### 2.12. Data analysis

Statistical analyses were conducted using GraphPad Prism v. 8.0.2 (GraphPad Software, San Diego, California USA). Outlier identification was performed with the ROUT method. Normality was evaluated using the Shapiro-Wilk test before performing statistical analyses. The statistical test and the number of independent replicates (n) used for each analysis are indicated in the respective figure legend. All data is presented as single data points and mean ± standard error of the mean (SEM). Differences were considered to be significant when p<0.05, being *p < 0.05; **p < 0.01; ***p < 0.001; ns: not significant. For data plotting, GraphPad Prism was employed. All data was collected and processed randomly.

## 3. RESULTS

### 3.1. ND523 microglial repopulation improves cognitive deficits induced by hTau^P301L^ overexpression in mice

Microglial repopulation has been shown to improve cognitive deficits in the short term (1 month) ^24^, but it remains unclear whether these effects persist over time. To address this, we evaluated its long-term impact in a tauopathy-induced model and explored whether the presence of an Nrf2 inducer (ND523) could help sustain or extend these benefits. We started by validating the CSF1-R inhibitor’s depletion efficacy treating adult male C57BL/6J mice with PLX5622 formulated in chow over a 14-days period - time window subsequently employed for microglial depletion. We confirmed that PLX5622 treatment produced a robust reduction, with an approximate 80-85 % decrease in microglia numbers compared to controls (Fig. S1, A-E), as previously reported ^14,24^.

Once validated microglial depletion, we proceeded with our experimental protocol. Mice were stereotaxically injected into the hippocampus, a crucial region for memory and learning highly affected in AD. They received either control AAV-GFP (Ctrl group) or AAV-hTauP301L. (Fig. 1A). P301L mutation represents one of the most common genetic variants of *MAPT* commonly used to study tau’s role in neurodegeneration ^1,40^. hTau-injected animals were then subdivided into four treatment cohorts: one receiving no further treatment (hTau group), one receiving PLX5622 followed by spontaneous repopulation (Repop group), one receiving PLX5622 followed by daily ND523-jelly administration (ND-Repop group), and one receiving only ND523-jelly administration (ND group) (Fig. 1B). Microglial numbers were assessed across these different conditions at the final point, and no significant differences were observed, indicating that microglial populations had returned to baseline levels after repopulation in all groups (Fig. S1, F-H).

**Figure 1.**
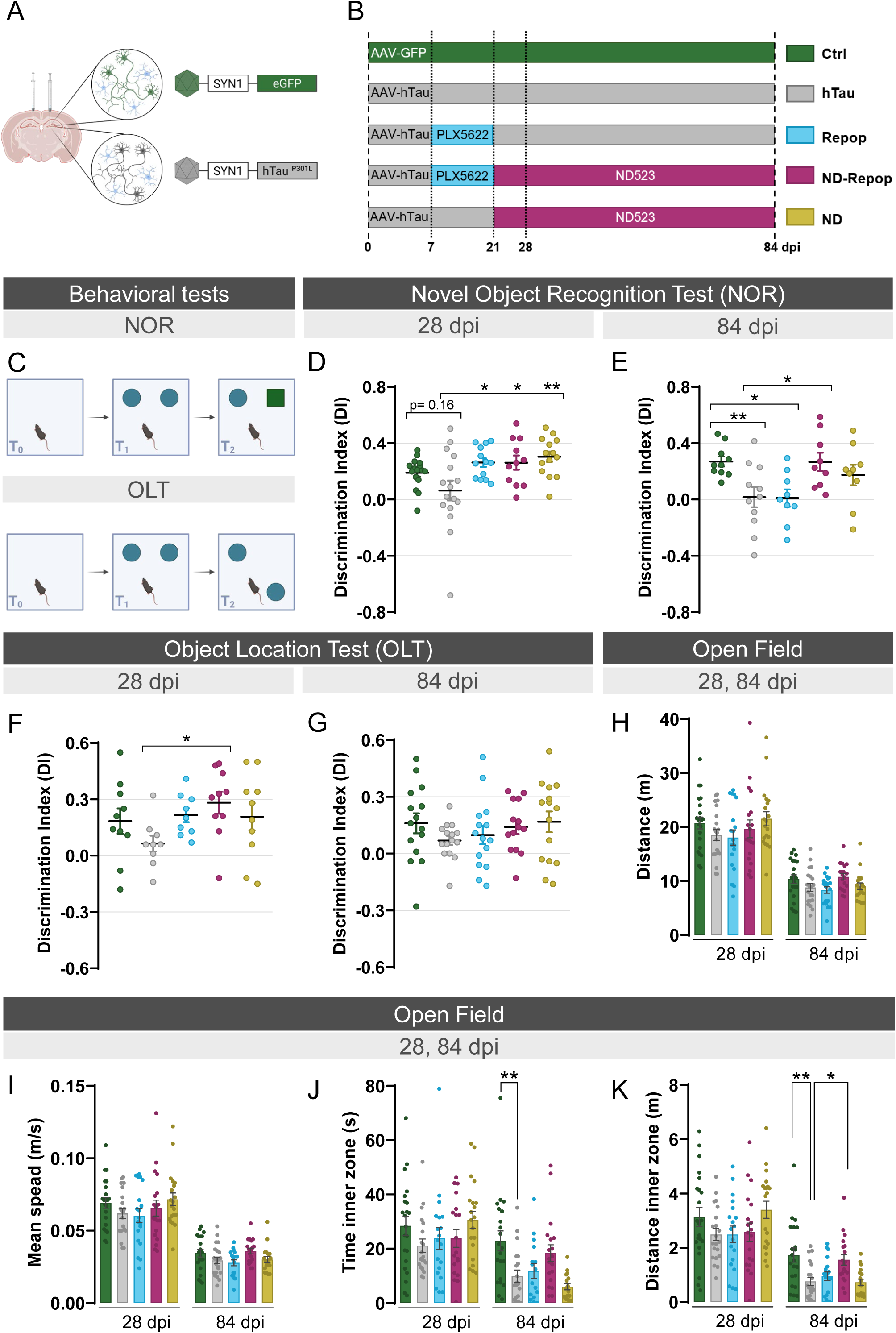
Microglial repopulation in the presence of ND523 improves cognitive deficits and behavioral alterations in a tauopathy-induced mouse model. **(A)** Schematic representation of the AAV-GFP (Ctrl)/AAV-hTau (hTau) bihippocampal stereotaxic induced model. **(B)** Experimental design illustrating the different groups and treatments. The Ctrl group received AAV-GFP injections, while the rest of the groups were injected with AAV-hTau and subdivided into non-treated (hTau), treated with PLX5622 (Repop), with PLX5622 followed by ND523 administration (ND-Repop) or with ND523 (ND). **(C)** Schematic representation of NOR and OLT tests. **(D,E)** DI in the NOR test 28 dpi (n = 13-16) and 84 dpi (n = 9-10 animals per condition). **(F, G)** DI in the OLT test 28 dpi (n = 9-10) and 84 dpi (n = 14-15 animals per condition). Open field analysis of **(H)** total traveled distance, **(I)** mean speed, **(J)** time spent in the inner zone and (K) distance travelled in the inner zone (n = 18-22 animals per condition). Data are represented as mean ± SEM. Significance was determined by one-way ANOVA followed by Tukey’s multiple comparisons test. Statistical significance indicated as *p < 0.05; **p < 0.01. Discrimination index (DI); novel object recognition (NOR); object location task (OLT); days post-injection (dpi). See also Fig S2.

Behavioral and cognitive assessments were conducted at two different time points, 28 and 84 dpi, to evaluate the impact of microglial repopulation on memory function over tau pathology progression. First, we performed the NOR test to evaluate recognition memory (Fig. 1C). At 28 dpi, hTau animals experience a slight memory impairment, represented by a decline in the DI, reaching values close to 0, whereas the rest of the hTau groups with the different treatments showed a normal cognitive state (Fig. 1D). However, at 84 dpi, when the repopulation window was extended in time, we observed a significant cognitive decline in the repopulation group, comparable to hTau animals. Interestingly, the combination of repopulation in the presence of ND523 prevented cognitive decline with a DI comparable to the control group (Fig. 1E). Spatial memory assessed by the OLT presented similar tendencies to those seen in the NOR test at both temporalities but it only showed a significant improvement for the the ND-Repop group at 28 dpi (Fig. 1F, G). Lastly, in the open field test, no differences in spontaneous motor activity were observed among the groups at any of the time points (Fig. 1H, I, Fig. S2). Interestingly, at 84 dpi, hTau animals exhibited anxiety-associated behavior, spending significantly less time and covering less distance in the inner zone of the box. This effect was notably reversed in the ND-Repop group, particularly in terms of distance traveled (Fig. 1J, K,). Thus, microglial repopulation alone was not able to perform long-term lasting functions. However, microglial repopulation in the presence of the Nrf2 inducer ND523 prevented cognitive decline and anxiety-related behaviors in a tauopathy mouse model in the long run.

### 3.2. ND523-mediated microglial repopulation fails to mitigate tau pathology but restores neuronal survival in the hippocampus of hTau^P301L^ injected mice

To investigate the contribution of repopulated microglia to tau pathology, we assessed different pathological phosphorylation epitopes of tau across relevant brain regions. Our tauopathy-induced mouse model exhibits rapid tau expression and phosphorylation ^41^. We first evaluated phosphorylation levels at Ser396/404 with the PHF1 antibody at the DG—the injection site—where hTau-injected mice displayed significantly increased levels compared to controls (Fig. 2A, D). However, none of the treatment conditions led to a significant reduction in the phosphorylation state at these residues. Next, we examined the phosphorylation at Ser202/Thr205 at the different hippocampal subregions through the AT8 antibody, which serves as the framework for Braak staging in AD patients ^42^. Consistent with our findings using PHF1 antibody, AT8 antibody revealed a significant increase in tau phosphorylation levels in hTau-injected animals compared to their matching controls (Fig. 2B, E). No significant differences in the phosphorylation levels of this epitope were detected across all treatment conditions. Interestingly, a modest but significant decrease was observed in phosphorylated Ser396/404 tau at CA3 in the ND-Repop (Fig. 2F). Previous studies, including work from our laboratory, have demonstrated severe neuronal loss in the granule cell layer (GL) thickness of the DG upon hTauP301L injection ^41,43^. To elucidate whether microglial repopulation followed by ND523 could mitigate this loss in hTau-injected mice, we assessed the GL integrity using DAPI staining (Fig. 2C). As expected, hTau mice exhibited a marked reduction in thickness, indicative of substantial neuronal loss (Fig. 2G). Interestingly, microglial repopulation followed by ND523 partially rescued this phenotype, while Repop and ND groups showed no evident neuroprotection. These findings suggest that microglial repopulation does not exert a major impact on tau pathology, but it can alleviate neuronal loss in the presence of ND523 in a pure tauopathy setting.

**Figure 2.**
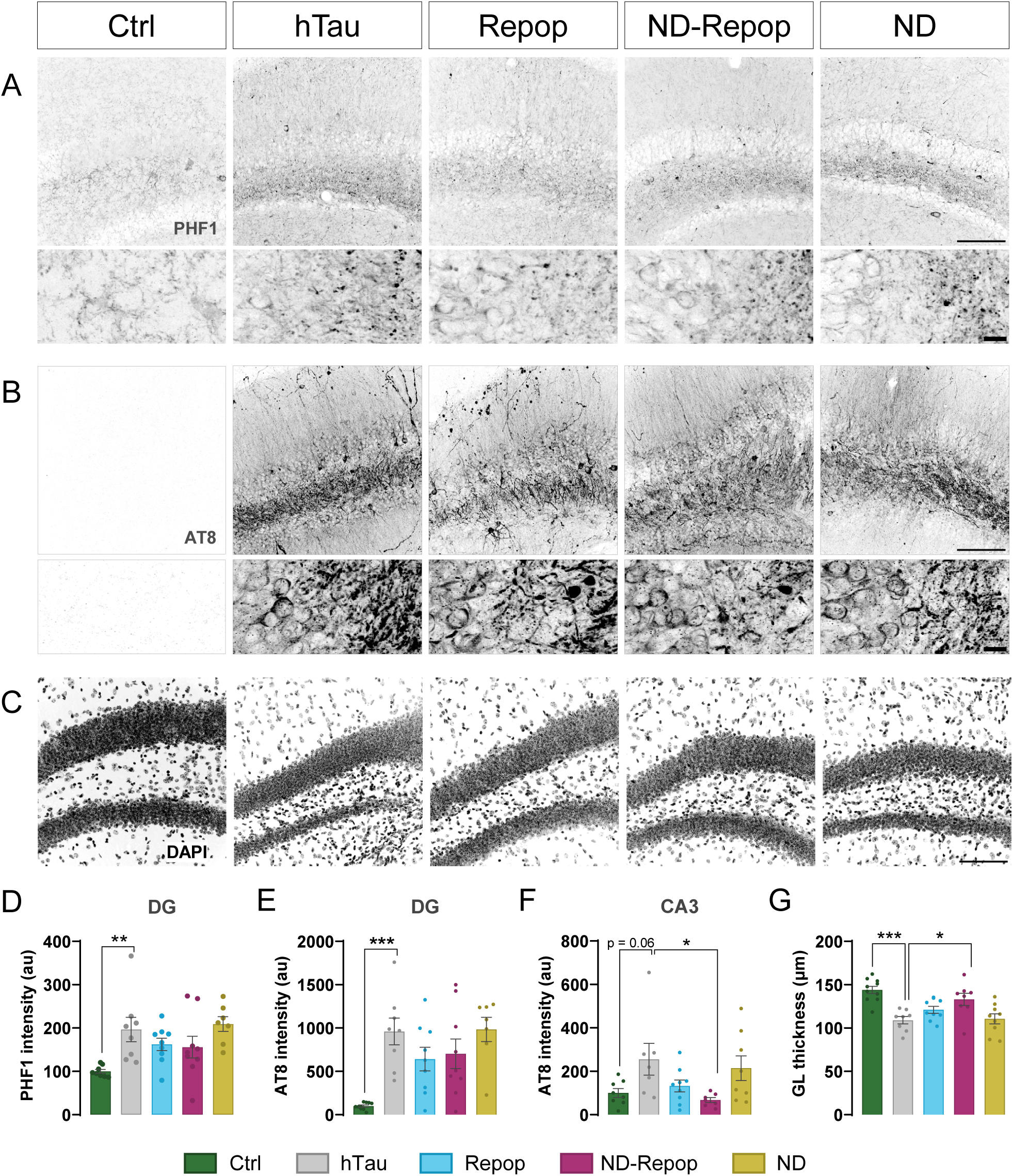
Microglial repopulation with ND523 fails to mitigate tau pathology but promotes neuronal survival in the hippocampus of hTau injected mice. Representative images of **(A)** PHF1 **(B)** AT8 and **(C)** DAPI immunostaining of the hippocampus. Quantification of **(D)** PHF1 in DG, **(E,F)** AT8 in DG and CA3, respectively, and **(G)** thickness of GL (n = 6-9 animals per condition). Significance was determined by one-way ANOVA followed by Tukey’s multiple comparisons test. Statistical significance indicated as *p < 0.05; **p < 0.01; ***p < 0.001. Scale bars: 100 µm and 10 µm in zoom images. Cornu ammonis 3 (CA3); days post-injection (dpi); dentate gyrus (DG); granulate layer (GL).

### 3.3. Microglial replacement followed by ND523 treatment mitigates microglial reactivity in a hTau^P301L^ mouse model

To gain insights into the temporal progression of microglial activity during repopulation, we assessed the microglial lysosomal marker Cd68, indicative of reactive microglia ^44^, at the intermediate and final timepoints; 42 and 84 dpi, respectively. At 42 dpi, hTau animals showed an increase in the intensity of Cd68 within Iba1^+^ cells compared to controls. At 84 dpi, microglial Cd68 levels continued to rise in all hTau-injected animals, except in the ND-Repop group, where this increase was halted (Fig. 3A, B). We have previously observed that the thickness of the DG is compromised upon human tau P301L overexpression. Interestingly, associations between this parameter and the levels of Cd68 showed a significant negative correlation (r= -0.60, p<0.0001), suggesting that sustained non-resolved microglial reactivity correlates with neurodegeneration (Fig. 3C). Since microglia morphology is a reliable indicator of cell functional state ^45^, we next performed morphological evaluation of these cells by segmentation analysis at 84 dpi (Fig. 3D). The parameters of area, perimeter, ramification index and Feret diameter were significantly decreased in hTau animals compared to control (Fig. 3E-G), indicative of a reduced morphological complexity (swollen cells with reduced branching) and a shift towards a more reactive state. Interestingly, only the ND-Repop group reverted these morphological changes, maintaining a more ramified morphology. Taken together, these results indicate that tau pathology drives microglia into a reactive state, with increased Cd68 levels and an ameboid-like morphology. These changes persist or even intensify over time, and spontaneous microglial repopulation does not seem to be able to stop them. However, microglial repopulation in the presence of ND523 mitigates these effects, effectively slowing down the progression of microglial reactivity.

**Figure 3.**
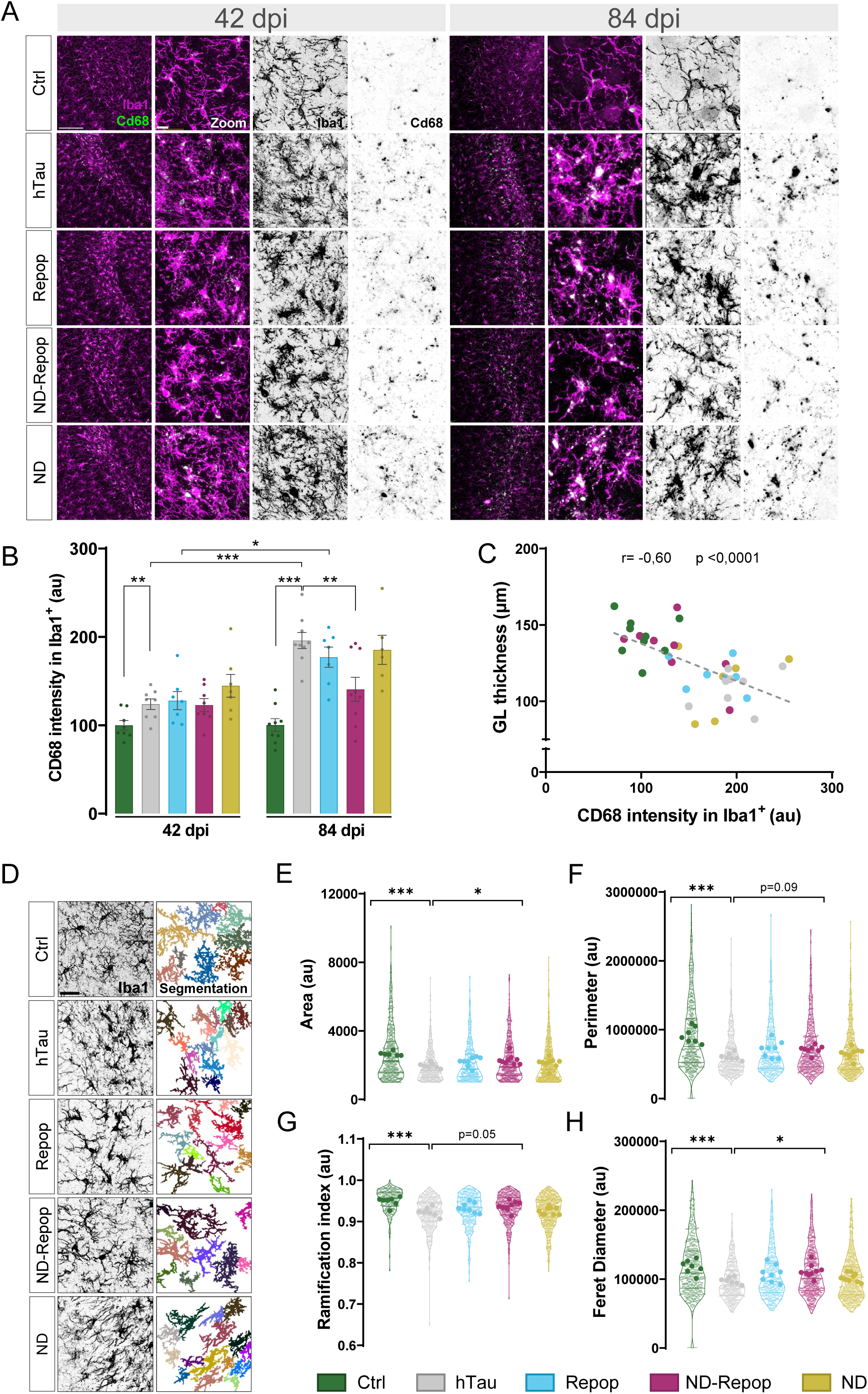
Long-term microglial repopulation in the presence of ND523 decreases Cd68 lysosomal marker and partially restores microglial morphology in hTau injected mice. **(A)** Representative images of Iba1 (magenta) and Cd68 (green) immunostainings of the hippocampus with a zoomed-in view alongside. **(B)** Quantification of Cd68 intensity within Iba1^+^ staining at 42 dpi and 84dpi (n = 6-9 animals per condition). **(C)** Correlation between DG layer thickness (reported in Fig. 2) and Cd68 intensity within Iba1 (n = 7-9). **(D)** Representative images of the segmentation analysis of Iba1 staining in brain hippocampal sections. Quantifications of **(E)** microglial area, **(F)** perimeter, **(G)** ramification index and **(H)** ferret diameter. An average of 60-70 cells per animal were analyzed. Small transparent dots represent all the individual cells analyzed and large colored dots represent the mean of each animal (n=7-9 animals per condition). Significance was determined by one-way ANOVA followed by Tukey’s multiple comparisons test. Pearson’s correlation was used to assess linear associations where indicated. Statistical significance indicated as *p < 0.05; **p < 0.01; ***p < 0.001. Scale bar: 100 µm and 10 µm in zoom images. Days post-injection (dpi); dentate gyrus (DG); granulate layer (GL). See also Fig S3.

### 3.4. Astrocyte reactivity exhibits a modest but non-significant decline after microglial repopulation followed by ND523 treatment in hTau^P301L^ injected mice

Microglia and astrocytes are closely intertwined, with their crosstalk being essential to maintain CNS homeostasis in both health and disease ^46^. Reactive microglia have been shown to drive astrocytes into a neurotoxic reactive state ^47^, potentially contributing to neuroinflammation and disease progression. Given this interaction and the presence of reactive microglia in our hTau-injected model, we sought to assess astrocyte reactivity. To do so, we performed double immunofluorescence staining for C3 and GFAP to better characterize astrocyte activation in this context (Fig. 4A). At 84 dpi, we observed a significant increase in GFAP mean intensity (Fig. 4B) and, most interesting, a significant increase in C3 intensity within GFAP^+^ cells (Fig. 4C) in hTau animals compared to their matching controls. Although this increase was not reverted in any experimental group, we did observe a strong trend towards reduced C3 reactivity in the ND-Repop group (p=0.08). These results are in line with those observed for microglial reactivity (Fig. 3), suggesting an interplay between microglia and astrocytes. In fact, when we performed a correlation analysis between C3 intensity within GFAP^+^ cells and Cd68 intensity within Iba1^+^ cells in the DG, we observed a positive and significant correlation between astrocyte and microglial reactivity (Fig. 4D, r= 0.61, p<0.0001). Interestingly, as previously observed for microglia (Fig. 3C), we also observed a significant negative correlation between astrocyte reactivity (measured as C3 intensity within GFAP^+^ cells) and GL thickness (Fig. 4E, r= -0.70, p<0.0001). These results suggest that modulating microglial activity through repopulation with an Nrf2 inducer may exert broader and more long-lasting effects than spontaneous repopulation alone, indirectly dampening astrocyte reactivity.

**Figure 4.**
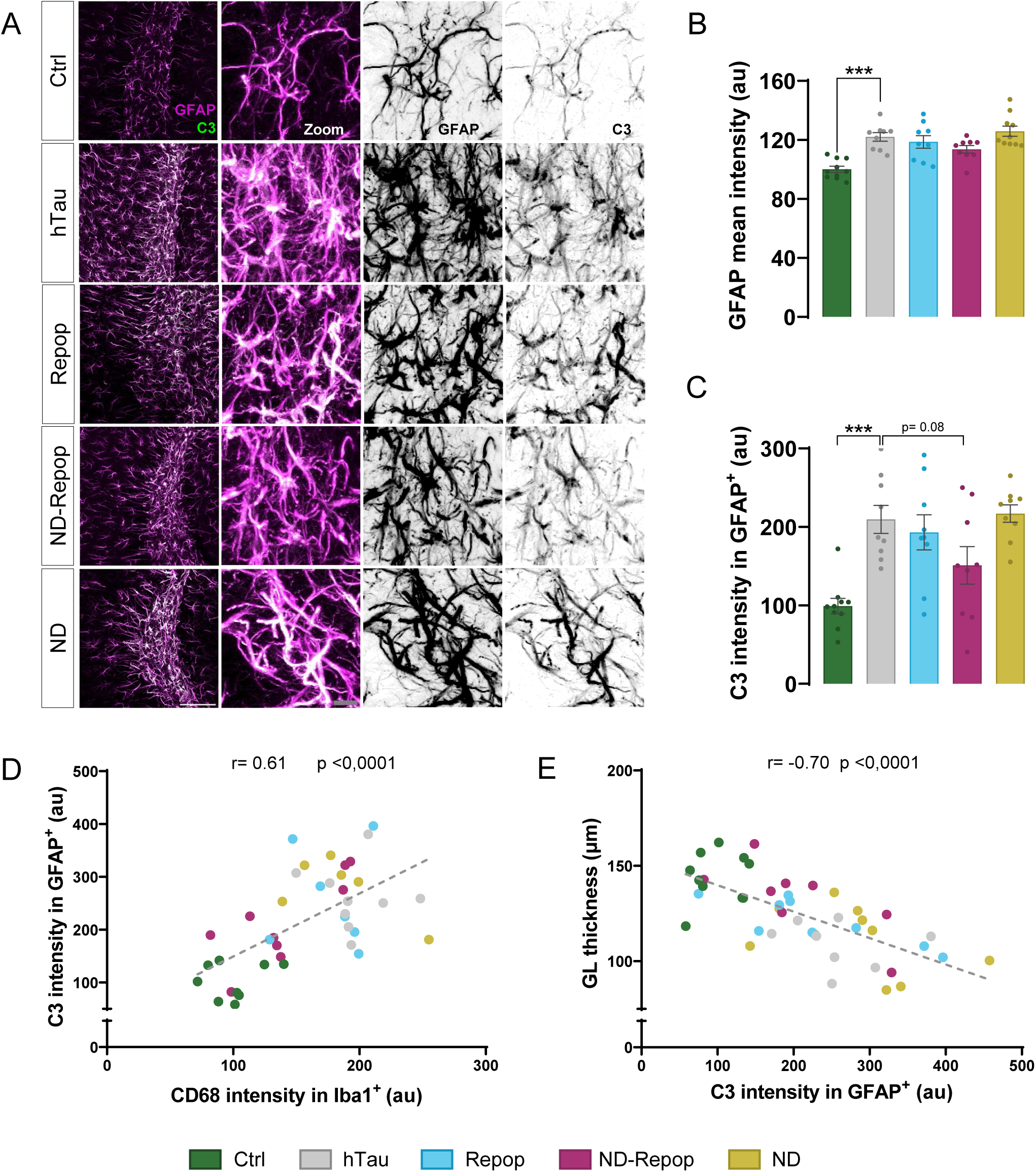
Astrocyte reactivity is not significantly reverted upon microglial repopulation followed by ND523 treatment in hTau-injected mice. **(A)** Representative images of GFAP (magenta) and C3 (green) immunostaining in the hippocampus with a zoomed-in view alongside. Quantification of **(B)** GFAP mean intensity and **(C)** C3 intensity within GFAP+ staining at 84 dpi (n = 9-10 animals per condition). Correlation analysis between **(D)** C3 intensity within GFAP and Cd68 intensity within Iba1 (reported in Fig 3) and **(E)** DG layer thickness (reported in Fig 2) and C3 intensity within GFAP (n = 7-10 animals per condition). Significance was determined by one-way ANOVA followed by Tukey’s multiple comparisons test. Pearson’s correlation was used to assess linear associations where indicated. Statistical significance indicated as *p < 0.05; **p < 0.01; ***p < 0.001. Scale bar: 100 µm and 10 µm in zoom images. Days post-injection (dpi); dentate gyrus (DG); granulate layer (GL).

## 3.5. ND523-mediated microglial repopulation modulates mitochondrial transcriptomic changes in hTau-injected mice

To investigate the molecular pathways underlying the benefits of the Nrf2-mediated repopulation, we performed bulk RNA sequencing of the hippocampus from Ctrl, hTau and ND-Repop mice at 84 dpi. After regressing out covariates, PCA revealed clear separation among the experimental groups (Fig. 5A). Differential gene expression analysis showed that hTau lesion alone showed a modest effect compared to Ctrl at this time point, resulting in only 10 DEGs (Supplemental Table 3)., while Nrf2 microglial repopulation showed a much stronger transcriptomic dysregulation, yielding 1,610 DEGs when compared to hTau (Fig. 5B, S4 and Supplemental Table 4).

**Figure 5.**
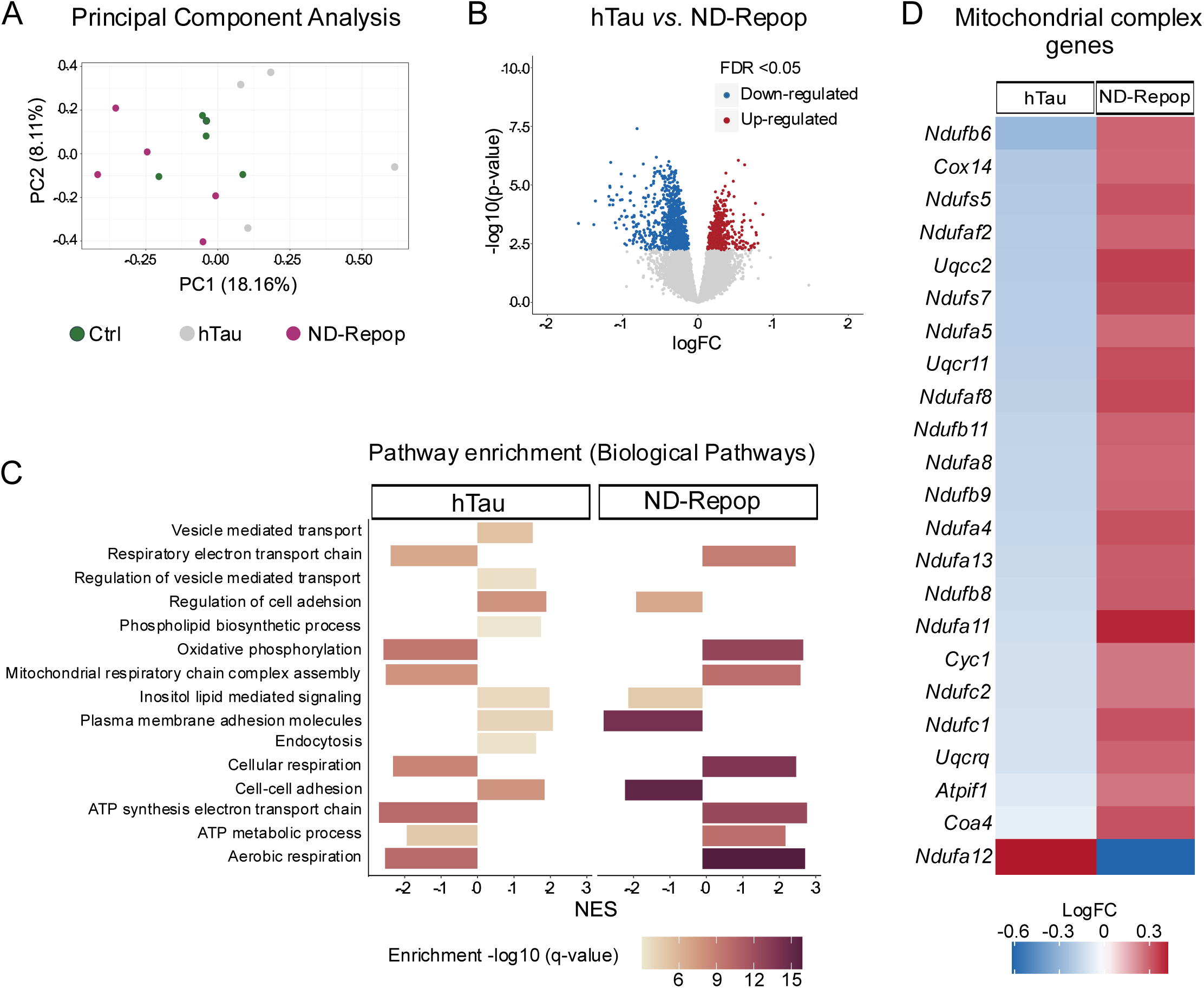
Transcriptional analysis of ND523-repopulated hippocampi reveals restoration of mitochondrial function impairments in tauopathy-induced mice. **(A)** PCA showing each RNA-seq sample colored by experimental group. **(B)** Volcano plot depicting log fold changes and -log10 of the p-value showing DEGs between hTau *vs.* ND-Repop mice. Each dot represents a specific gene; blue genes have an FDR>0.05 and red genes have an FDR<0.05. **(C)** Pathway enrichment analysis using GO Hallmark Pathways and a pre-ranked GSEA. Color key corresponds to the -log10 of the q-value for enrichment score. **(D)** Mitochondrial complex genes expression. Color key corresponds to the log fold change (n = 4-5 animals per condition). Differentially expressed genes (DEGs); false discovery rate (FDR); gene ontology (GO); gene set enrichment analysis (GSEA); principal component analysis (PCA).

To explore the biological significance of these transcriptional changes, we conducted GSEA focusing on biological pathways. Numerous pathways were significantly enriched in both the hTau and ND-Repop conditions (Supplemental Table 5). A representative subset is shown in Fig. 5C, where hTau injection was associated with impaired mitochondrial performance, with a downregulation of mitochondrial respiration and ATP synthesis pathways, alongside an upregulation of pathways involved in lipid metabolism, vesicle-mediated transport, cell adhesion, and endocytosis. Notably, some of these transcriptional changes were reversed in ND-Repop animals, with significant upregulation of mitochondrial respiration pathways and downregulation of those related to lipid metabolism and cell adhesion.

Given the robust effect on mitochondrial function, we focused on the expression of genes encoding for mitochondrial complex components (Fig. 5D). hTau injection resulted in reduced expression of many mitochondrial-related genes, whereas Nrf2-mediated microglial repopulation significantly restored their expression. Overall, our data points to mitochondrial homeostasis modulation by ND523 as a key mechanism underlying its sustained benefit on microglial repopulation.

## 4. DISCUSSION

Epidemiological, clinical, and experimental research has established that innate immune activation plays a critical role in the pathogenesis and progression of AD ^48^. Therefore, strategies aimed at resetting the immune system by depleting these cells and letting them spontaneously repopulate have held great promise for AD and other neurodegenerative diseases ^49^. Although this approach has shown positive results in the short-term and, mostly, in Aβ models ^25,50,51^, whether these effects are sustained over time or need to be pharmacologically modulated to be further extended remains largely unknown. Here, we have defined the potential of long-term microglia repopulation, implementing a spontaneous repopulation alone or in combination with an Nrf2 inducing compound in a pure tauopathy model.

Our study was conducted using an induced h-TauP301L overexpressing mouse model that exhibits key pathological features of AD, including tau pathology, gliosis, neuronal loss, and cognitive impairment. This model offers the advantage of a shorter disease progression timeline compared to traditional AD models, where tau pathology develops over months or even years ^52–55^. Notably, it exhibits sustained microglial reactivity, allowing us to assess how microglial reprograming impacts on disease progression. To ensure microglial elimination and subsequent repopulation, we employed PLX5622, which effectively depleted around 85 % of microglia, similarly to other studies ^14–17,56,57^. Microglia numbers have been reported to be restored within 3 weeks following inhibitor withdrawal ^18,58,59^. In line with this, our results showed that microglia numbers among PLX-depleted groups were broadly similar and mirrored those of control animals after the repopulation phase (Fig. S1, F-H). The origin of these newly repopulated cells has been widely debated. However, recent findings increasingly support that they arise from proliferation of surviving microglia, ultimately restoring microglial density to that of untreated controls ^17^. This remarkable capacity for regeneration provides a valuable framework for therapeutic studies aimed at resetting dysfunctional microglia in neurodegeneration.

Our results showed that spontaneous microglial repopulation had initial benefits on memory measured in the NOR test. However, these effects were lost over time. In contrast, when repopulated microglia were pharmacologically modulated with ND523, the cognitive improvements were maintained over time. Even though similar tendencies appeared in the OLT, the differences in test sensitivity to hippocampal-dependent deficits and the added variability from spatial navigation complexity might explain the lack of significance and dispersion observed. Although behavioral studies on microglial repopulation are scarce, similar improvements have been reported in spatial memory assessed by the Morris Water Maze in aged ^24^ and in 5xFAD mice ^25^ in the short-term. Locomotor activity was similar across all groups, indicating that alterations in movement did not confuse cognitive assessments results. However, there was a general decrease in locomotor parameters from 28 to 84 dpi, probably due to the loss of novelty in the testing environment. Notably, only distance and time spent in the inner zone parameters were different between hTau-injected mice and controls at 84 dpi, indicative of an anxiety-associated behavior previously observed in other tauopathy models ^60–62^. Interestingly, this behavior was reversed in the ND-Repop. Therefore, pharmacological activation of Nrf2 signaling pathways during microglia repopulation mitigates cognitive impairment and anxiety-related symptoms in our tauopathy model.

We found that microglial repopulation, either spontaneous or in the presence of ND523, did not alter tau phosphorylation at the DG. These findings aligned with previous studies that also reported no reduction in AT8 levels when tau was seeded into 5xFAD mice brain after the repopulation period ^63^. Additionally, 3xTg old mice revealed varying outcomes depending on the specific tau phosphorylated epitope analyzed ^64^. Interestingly, AT8 levels were significantly reduced in CA3 upon ND523 microglial repopulation. This reduction may imply an attenuation in the migration patterns of the overexpressed tau protein from the DG to other hippocampal regions. This was in line with previous studies showing microglial implication in tau propagation through exosomes and how their depletion suppressed propagation in an AAV-hTau-based model ^20^. Considering the implication of the DG in memory retrieval ^65^, neurodegeneration within this region could help explain the results observed in the behavioral tests. In fact, hTau-injected animals suffered a significant neuronal loss which was significantly reduced following microglial repopulation with ND523. These results indicate that this combined strategy affords certain degree of neuroprotection in a tau-driven pathology. However, whether these effects translate into improved neuronal function or integrity warrants further investigation.

Given evidence that microglial repopulation can restore altered microglial phenotypes, we investigated whether this holds true in our tauopathy model. Our findings suggest that while initial reactivity was comparable among hTau-injected groups, long-term outcomes diverged, and ND523-treated repopulated microglia exhibited a more stable reactive profile and restored morphology. Notably, previous studies have reported microglial repopulation to restore microglial morphological phenotypes under different pathological conditions, including the 5xFAD model of amyloidosis ^25^, traumatic brain injury ^66^, and neurotoxic pre-primed microglia exposed to LPS ^67^. Similarly, repopulated microglia have been shown to normalize Cd68 expression in aged mice ^24^. However, other authors reported that, despite this normalization, these cells exhibited resistance to sustained phenotypic changes, still adopting a pro-inflammatory state ^68^. These results highlight the fact that the reversal of microglial phenotypes upon repopulation might be highly context-dependent, and discrepancies may stem from model aggressivity and chronicity of disease. Hence, the sustained and aggressive nature of our hTau overexpression model may hinder a phenotype reversal following repopulation. Thus, microglia may inherit a reactive profile due to persistent pathological cues in their environment early after repopulation, as hypothesized in ^24,69^. Notably, when repopulated microglia were pharmacologically modulated with ND523, both Cd68 reactivity and morphological restoration showed a clear effect. These findings suggest that spontaneous microglial repopulation alone is not sufficient to fully counteract tau-driven microglial response without any additional intervention and highlight Nrf2 as a potential therapeutic target for modulating repopulated microglia.

To address the impact of microglial repopulation in tau pathology, it is essential to consider the communication between microglia and astrocytes. These two cell types are known to communicate closely and shape each other’s responses in neurodegenerative conditions ^70^. Interestingly, previous studies on microglial repopulation have primarily focused on changes within microglia themselves, without thoroughly assessing the effects on other cell populations. Here, we assessed astrocyte reactivity through the analysis of C3 intensity. hTau-injected animals displayed increased C3 levels, indicating the presence of astrogliosis, which failed to be reduced with spontaneous repopulation alone. Similarly, ÒNeil *et al*. did not observe astrocyte phenotype change upon repopulation in aged mice and highlighted the importance of the brain environment in ultimately driving cell priming ^68^. This aligns with our results where newly repopulated ND523-shaped microglia partially reverted C3 reactivity. Hence, pharmacological modulation of repopulated cells seems to sustain their homeostatic phenotype, thereby positively influencing the cells they interact with. This highlights the importance of communication between glia, as well as with neurons in shaping disease progression and underscores the importance of studying cell populations in a more holistic manner.

Lastly, to better understand the molecular mechanisms underlying the beneficial effects of Nrf2 activation during microglia repopulation, we performed bulk RNA-seq analysis. Our data revealed significant downregulation of pathways central to mitochondrial energy metabolism driven by hTau-P301L overexpression. These alterations included a significant downregulation of oxidative phosphorylation components that might encompass disruptions in the electron transport chain and ATP synthesis in our tauopathy model. These alterations align with accumulating evidence implicating mitochondrial bioenergetics dysfunction as a central feature in AD pathogenesis ^71^. Tau interacts extensively with proteins involved in mitochondrial bioenergetics and P301L mutation has been shown to impair its energy metabolism by weakening interactions with subunits of the electron transport chain and ATP/ADP transporters in iPSC-neurons ^72^. This mechanism is further supported by proteomic analysis from mice carrying a similar mutation that showed reduced levels of respiratory chain proteins, and an age-dependent reduction in mitochondrial respiration and an impaired ATP production ^73^. Therefore, alterations in mitochondrial components seem to play a critical role in disease pathogenesis, and their decreased expression or weakened association with tau could contribute to mitochondrial dysfunction and impaired cellular energetics underlying neurodegeneration.

Remarkably, treatment with ND523 during the microglia repopulation phase restored mitochondrial energetic disturbances caused by hTau, further supporting the growing evidence of Nrf2 as a promising therapeutic target to counteract mitochondrial dysfunction ^74^. Nrf2 regulates redox homeostasis, mitochondrial membrane potential, and ATP synthesis. It also plays a critical role in mitochondrial fatty acid oxidation and helps maintain the structural and functional integrity of mitochondria ^75^. Consistent with this, the Nrf2 inducer sulforaphane has been shown to upregulate different subunits of the respiratory complexes and ATP synthase *in vivo* ^76^. Similarly, omaveloxolone, a clinically approved Nrf2 activator, enhanced mitochondrial substrate availability and activity, prevented from oxidative stress and decreased mitochondrial energy imbalance ^77,78^.

Our findings suggest that ND523-mediated Nrf2 induction during microglial repopulation improved their bioenergetic profile altered upon tau pathology. Concretely, microglia undergo metabolic reprogramming shifting from oxidative phosphorylation toward glycolysis in response to pathological stimuli (e.g. NFT) to rapidly support the heightened metabolic demands required for surveillance and phagocytosis ^79^. ND523 may support the energy state of new microglial cells, enabling them to maintain their functions and avoid pathological reactivity. Furthermore, the Nrf2-driven benefits may extend beyond microglia to other energy-demanding cells like neurons. Nonetheless, further research is required to clarify the effect of ND523 in Nrf2-mediated pathways in shaping repopulated microglial functional state.

## 5. CONCLUSIONS

While microglial repopulation alone may offer limited benefits in the context of tau pathology, its combination with the Nrf2 inducer ND523 confers neuroprotective effects and attenuates key pathological features: microglial reactivity, mitochondrial dysfunction, hippocampal neuronal loss and cognitive decline. Whether these beneficial effects persist for longer periods than 84 dpi requires further investigation. Nevertheless, these findings emphasize the importance of shaping the fate of self-renewed microglia and point pharmacologically controlled microglia repopulation, for instance via Nrf2 activation, as a potential therapeutic avenue to mitigate neurodegeneration in tauopathies.

## AVAILABILITY OF DATA AND MATERIALS

The datasets analyzed during the present study are available from the corresponding author on reasonable request.

## DECLARATIONS

All animal experiments were conducted according to the European Union Directive 2010/63/EU and the national Spanish Royal Decree for Animal Protection RD53/2013. They were authorized by the Institutional Ethics Committee of the Autonomous University of Madrid and the Community of Madrid, Spain (PROEX: 218.5/20).

## AUTHOR CONTRIBUTIONS

**LV:** Conceptualization, Methodology, Validation, Formal analysis, Investigation, Data Curation, Writing - Original Draft, visualization. **MGL:** Conceptualization, Methodology, Resources, Writing - Review & Editing, Supervision, Project administration and Funding acquisition. **ET:** Conceptualization, Methodology, Writing - Review & Editing, Supervision and Project administration. **EN:** Software, Validation, Formal Analysis, Visualization. **PN:** Investigation. **MIRF** and **JB:** Resources.

## DECLARATION OF GENERATIVE AI AND AI-ASSISTED TECHNOLOGIES IN THE WRITING PROCESS

During the preparation of this work the authors used ChatGPT (OpenAI) in order to improve language, clarity, and structure. After using this tool, the authors reviewed and edited the content as needed and take full responsibility for the content of the publication.

## ACKNOWLEDGEMENTS

We would like to thank María Dolores Vallejo from the Confocal Unit of the Universidad Autónoma de Madrid for its support during image acquisition and David Muñoz from the Animal Facilities of the Universidad Autónoma de Madrid.

## SUPPLEMENTARY FIGURES

**Figure S1. Microglial depletion after PLX5622 administration for 14 days in chow. (A)** Representative images of the hippocampus and cortex stained with Iba1 (green)/ DAPI (blue) and PU.1 (green)/ DAPI (blue) under control or PLX5622 conditions. Quantification of the number of **(B)** Iba1^+^ and **(C)** PU.1^+^ cells in the hippocampus (n=3 animals per condition). Quantification of the number of **(D)** Iba1^+^ and **(E)** PU.1^+^cells in the cortex (n=3). **(F)** Representative images of the hippocampus and cortex stained with PU.1 (red) under the different experimental conditions studied. Quantification of the number of PU.1^+^ cells in the **(G)** hippocampus and **(H)** cortex (n=8-9 animals per condition). Scale bar: 100 µm (top) and 50 µm (bottom). Significance was determined by un-paired t-test test and one-way ANOVA followed by Tukey’s multiple comparisons. Statistical significance indicated as *p < 0.05; **p < 0.01; ***p < 0.001; ****p < 0.0001. Cortex (CTX); Hippocampus (HP).

**Figure S2. Parameters assessed in the open field test at 28 and 84 dpi. (A)** Maximum speed, **(B)** time spent in the outer zone, **(C)** distance travelled in the outer zone, **(D)** line crossings, **(E)** time mobile and **(F)** time immobile (n = 18-22 animals per condition). Data are represented as mean ± SEM. Significance was determined by one-way ANOVA followed by Tukey’s multiple comparisons test. Statistical significance indicated as *p < 0.05; **p < 0.01. Days post-injection (dpi).

**Figure S3. Microglial morphology analyzed by segmentation analysis at 84 dpi. (A)** Average thickness and **(B)** geodesic diameter. An average of 60-70 microglial cells per animal were analyzed. Small transparent dots represent all the individual cells analyzed and large colored dots represent the mean of each animal (n=7-9 animals per condition). Significance was determined by one-way ANOVA followed by Tukey’s multiple comparisons test. Statistical significance indicated as *p < 0.05; **p < 0.01; ***p < 0.001.

**Figure S4. DEGs in the hippocampus of hTau-injected animals compared to control at 84 dpi. (A)** Volcano plot depicting log fold changes and -log10 of the p-value showing DEGs between hTau *vs.* Ctrl mice. Each dot represents a specific gene; blue genes have an FDR>0.05 and red genes have an FDR<0.05 (n = 4-5 t animals per condition). Differentially expressed genes (DEGs); false discovery rate (FDR).

