## Supplementary figures and images for "Long-term Nrf2-driven microglial repopulation mitigates microgliosis, neuronal loss and cognitive deficits in tauopathy"

### Supplementary figures

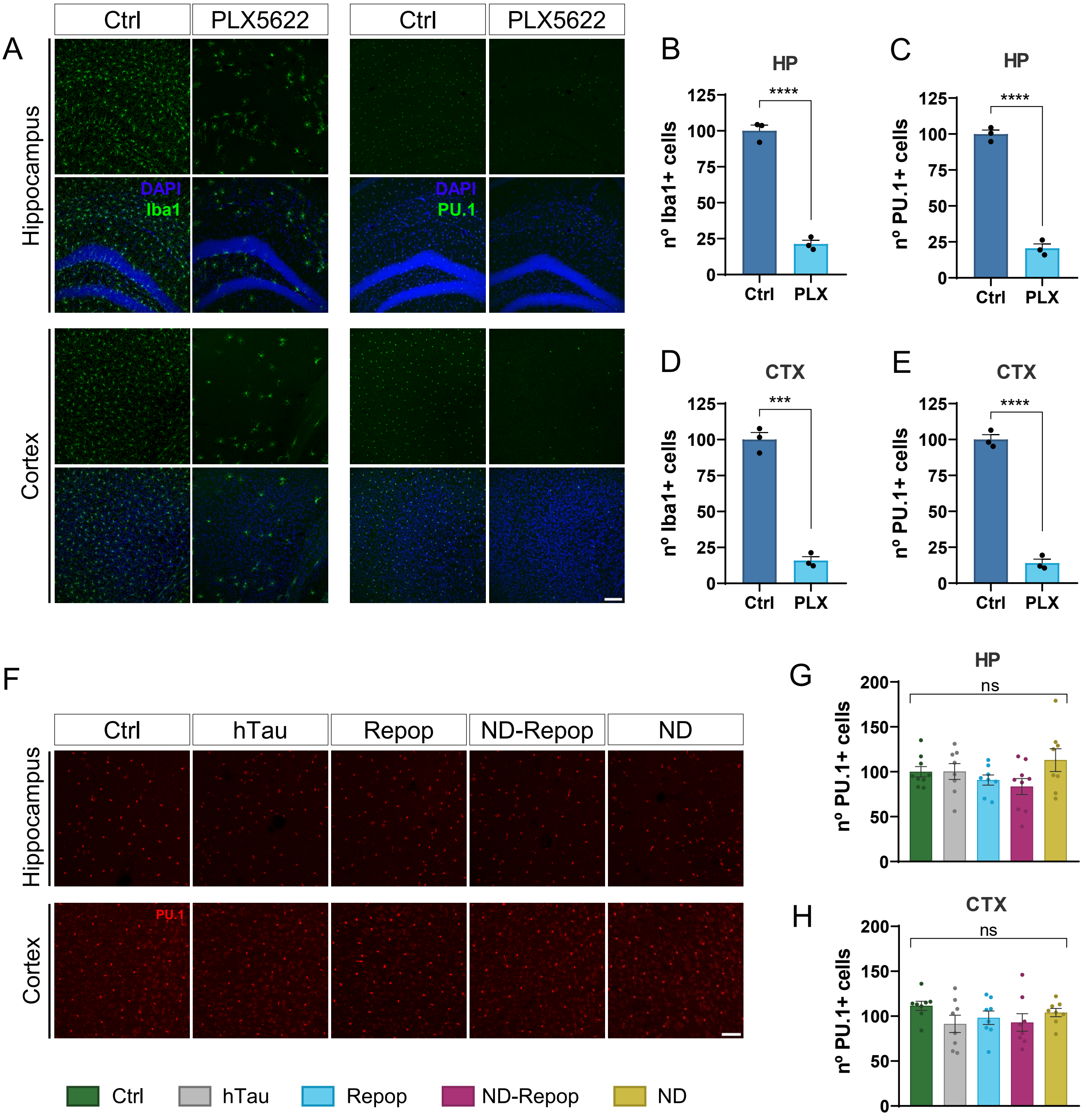

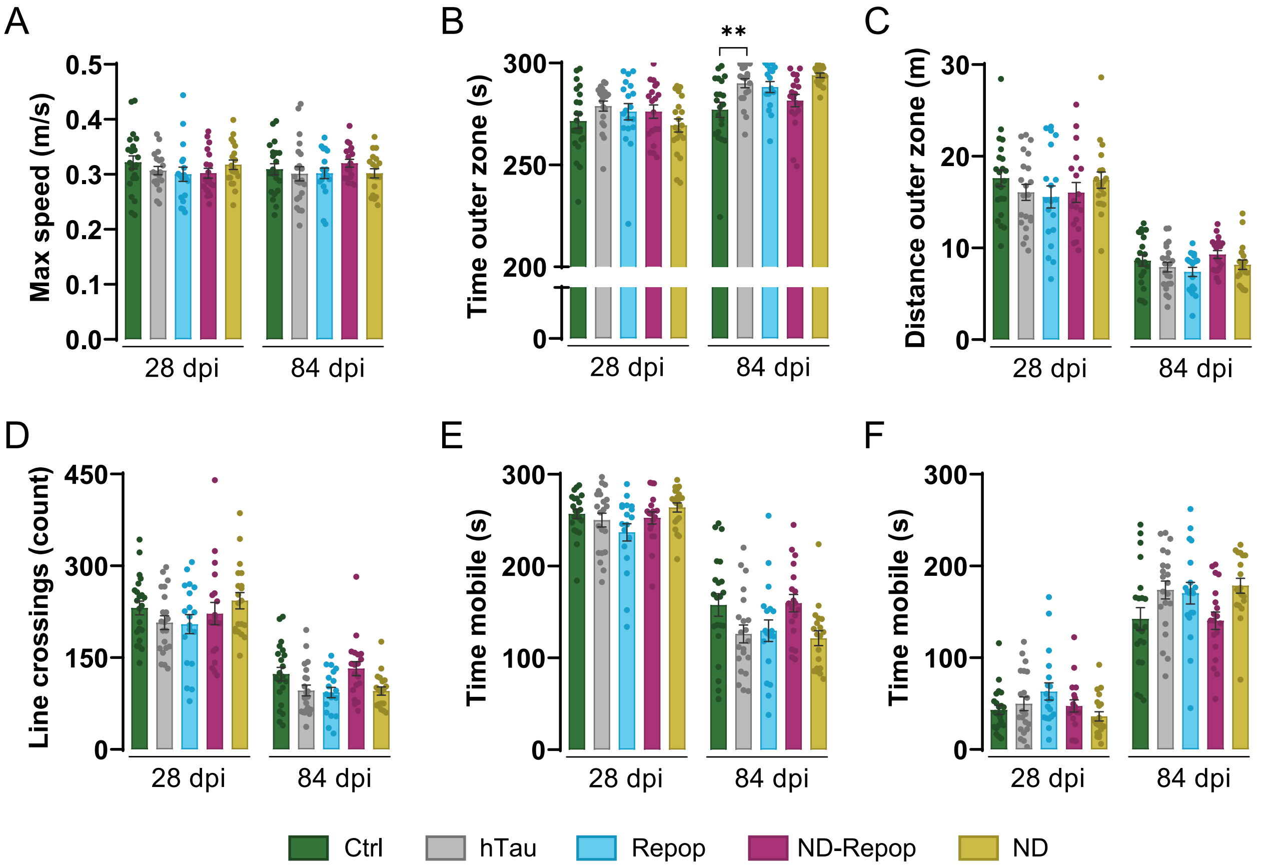

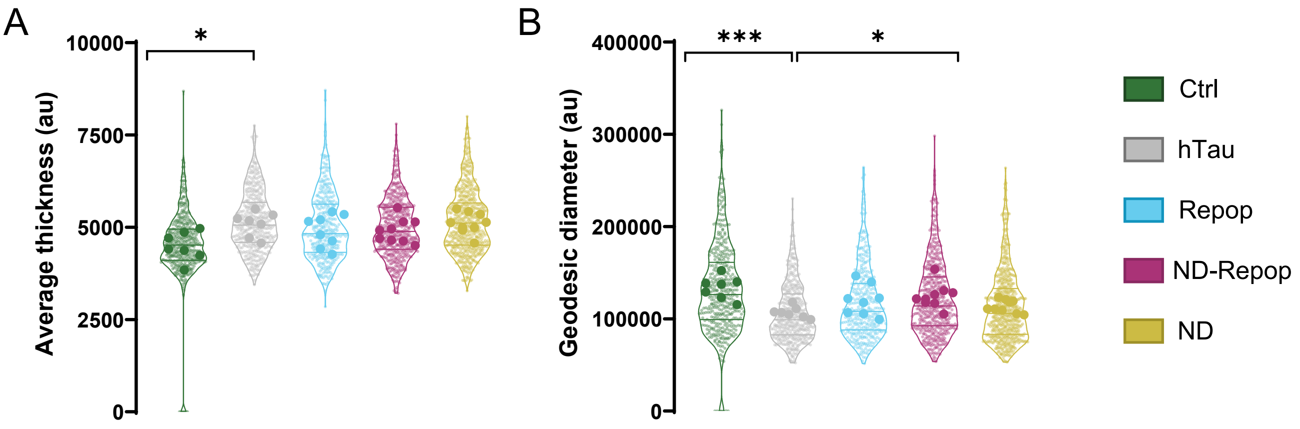

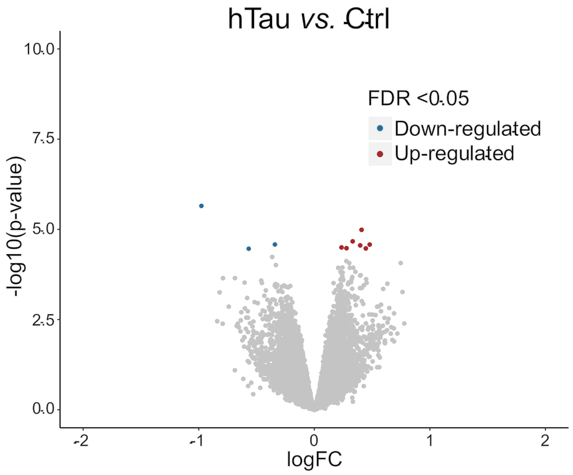
